# ABACUS: A flexible UMI counter that leverages intronic reads for single-nucleus RNAseq analysis

**DOI:** 10.1101/2020.11.13.381624

**Authors:** Simon Xi, Lauren Gibilisco, Markus Kummer, Knut Biber, Astrid Wachter, Maya Woodbury

**Author notes:** Co-senior authors (Astrid Wachter, Maya Woodbury). Corresponding author: Simon Xi.

## Abstract

Single-nucleus RNA sequencing (sNuc-RNAseq) is an emerging powerful genomics technology that combines droplet microfluidics with next-generation sequencing to interrogate transcriptome changes at single nucleus resolution. Here we developed Abacus, a flexible UMI counter software for sNuc-RNAseq analysis. Abacus draws extra information from sequencing reads mapped to introns of pre-mRNAs (~60% of total data) that are ignored by many single-cell RNAseq analysis pipelines. When applied to our pilot human brain sNuc-RNAseq data, ABACUS nearly doubled the number of nuclei identified by the CellRanger workflow, recovering a large number of nuclei from non-neuronal cells. By incorporating intronic reads into gene expression quantification, we showed that they encoded additional and valid transcription features of individual cells and could be used to improve cluster resolution of different cell types. By separately counting UMIs derived from forward and reverse intronic reads and from exonic reads, Abacus gives users flexibility in representing genes expressed at different abundance levels. In summary, Abacus represents a flexible, improved workflow for sNuc-RNAseq data processing and analysis.

## Introduction

Recent advances in single-cell RNAseq technologies have enabled large-scale profiling of cellular heterogeneity at unprecedented resolution. While most single-cell RNA sequencing methods require fresh tissues for high-quality single cell preparation, single-nucleus-RNAseq (sNuc-RNAseq) has the advantage of being applicable to frozen and archived tissue samples that are more readily available. sNuc-RNAseq has rapidly become the method of choice to study cell-type specific gene expression in human postmortem tissues under different development stages and disease conditions (Habib et al. 2017; 2016; Mathys et al. 2019; Kim et al. 2018). Similar to single-cell methods, droplet-microfluidics based sNuc-RNAseq techniques such as DroNc-seq (Habib et al. 2017) and the 10x Genomics Chromium system capture single nuclei in individual oil droplets. Then transcripts in each nucleus within each droplet are reverse transcribed with oligo-dT primers that each contain a common barcode sequence and a unique molecular identifier (UMI) sequence. These cDNA products are pooled, PCR amplified, and then sequenced. Barcode sequences are used to assign sequencing reads to individual nuclei, while UMI counts provide gene expression estimates in each nucleus. For data analysis, pipelines such as CellRanger (Zheng et al, 2017) originally designed to analyze single-cell data are commonly used. However, nuclei differ substantially from cells in their RNA contents. Only a small fraction of total mRNAs in the cells reside in the nucleus, but pre-mRNAs with intronic sequences are primarily found in the nucleus. It has been previously observed that the majority of sequencing reads generated in sNuc-RNAseq studies map to intronic regions of the genome. Most analysis pipelines originally developed for single-cell RNAseq studies only make use of reads derived from exons. Some pipelines provide an option to combine the UMI counts from intronic and exonic reads as gene expression estimates without a thorough justification, e.g. CellRanger (Zheng et al, 2017). To help alleviate this data analysis gap, we developed a software package called Abacus that provides a flexible workflow to derive UMI counts from exonic and intronic reads in both the forward and reverse directions (i.e. in the same or opposite direction of the mRNA). When applied to a sNuc-RNAseq dataset of human postmortem brain, we further demonstrated the utility of Abacus-derived UMI counts to improve the identification of nuclear barcodes, cluster analysis, and gene expression quantification.

## Implementation

### Nuclei isolation and single nuclei library preparation

Nuclei isolation was performed using ~300mg of frozen brain tissue from the cortex of a non-diseased human donor. The tissue had a RIN value of 6.4. Briefly, tissue was homogenized in 4mL of lysis buffer (0.32M Sucrose, 5mM CaCl_2_, 3mM Mg(Ac)_2_, 0.1mM EDTA, 10mM Tris-HCl, pH 8, all sterile and DNAse/RNAse free, with 0.1% triton x-100, and 1mM dithiothreitol (DTT) in sterile H_2_0) using a glass Wheaton homogenizer on ice. Lysate was layered on top of a sucrose solution (1.8M sucrose, 3mM Mg(Ac)_2_, 10mM Tris-HCl, pH8, and 1mM DTT in sterile H_2_0) in an ultracentrifuge tube (#244060, Beckman Coulter). Nuclei were separated from cellular debris by ultracentrifugation (107,000 x g for 1.5 h at 4°C). The supernatant was removed and pellets were resuspended in buffer containing DPBS, 0.2U/ul RNAse inhibitor (Roche) and 10ug/ml ultrapure BSA (Thermo Fisher). Nuclei were counted with EVE automated Cell Counter (Nanoentek). Nuclei concentration was adjusted according to the manufacturer’s instructions before single nuclei lysis and barcoding using the 10X Genomics Chromium controller. Once the uniquely-barcoded beads called GEMs (Gel Bead-In Emulsions) were collected after the Chromium controller runs, sequencing libraries were prepared using Chromium single cell 3’-reagent V3 kit following the vendor’s recommended protocol. Sequencing was performed on an Illumina HiSeq2500. For data QC, sequencing reads were processed using the standard CellRanger v3 pipeline. Basic QC metrics such as total read counts and number of reads per nucleus were taken from the standard CellRanger report.

### UMI counting with Abacus

Sequencing reads were first processed using the standard CellRanger pipeline. BAM files generated by CellRanger were then subjected to Abacus. The tool parses the BAM files and extracts the barcodes and corrected UMI sequences from aligned reads, then summarizes the UMI counts from intronic and exonic reads in the forward and reverse directions for each gene. Final UMI count tables (genes as rows and barcodes as columns) are generated in the sparse matrix format as the CellRanger output. In total, three UMI count tables are generated separately: one from forward exonic reads, one from forward intronic reads, and one from reverse intronic reads (see Figure 1).

**Figure 1.**
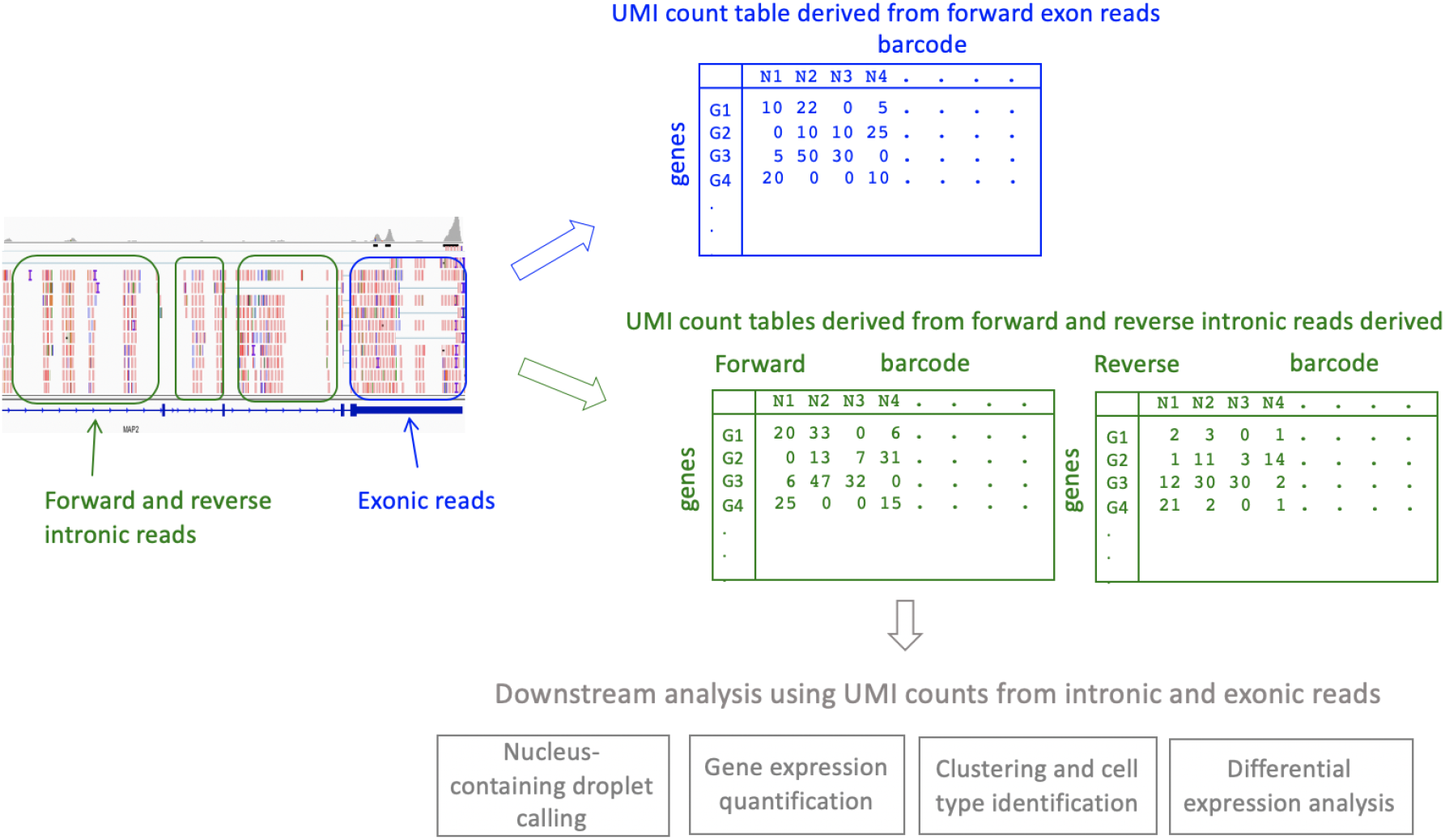
Schematic representation of the Abacus intronic and exonic read UMI counter workflow. Abacus parses CellRanger-derived BAM files and extracts the barcodes and corrected UMI sequences from aligned reads, then summarizes UMI counts from intronic and exonic reads in the forward and reverse directions for each gene. Final UMI count tables (genes as rows and barcodes as columns) are generated in sparse matrix format. In total, three UMI count tables are generated, from 1) forward exonic reads, 2) forward intronic reads, and 3) reverse intronic reads. These can then be used for downstream analyses such as clustering and differential expression.

### Downstream data analysis

To identify nuclear barcodes, Abacus first summarizes the counts in each UMI count table by column (i.e. by barcode). Afterwards, each barcode is associated with three UMI count categories, one from forward exonic reads, one from forward intronic reads, and one from reverse intronic reads. Abacus then filters nuclear barcodes by UMI counts from forward exonic reads and from forward intronic reads. The default filtering threshold is UMI count > 80. For clustering analysis, count tables generated by Abacus were loaded into R package v3.5 and underwent dimensionality reduction in “Seurat” package v2.3 (Butler et al. 2018) with default parameters. To run clustering analysis jointly using UMI counts from forward exonic reads and forward intronic reads, the UMI count table from forward intronic reads was appended to the UMI count table from forward exonic reads. The combined UMI count table was subjected to dimensionality reduction in “Seurat” package v2.3 (Butler et al. 2018).

## Results

### Different sources of intronic reads

It has been reported in sNuc-RNAseq studies that a large proportion of sequencing reads were aligned to intronic regions of the genome (Habib et al. 2017). It was suggested that these intronic reads were derived from mis-priming of the oligo-dT primers to the poly-A repeats in the intronic regions of the pre-mRNAs during the first-strand cDNA synthesis by reverse transcription (La Manno et al. 2018). However, in the nucleus, both pre-mRNAs and genomic DNA contain poly-A repeats that could potentially provide the template for oligo-dT priming and cDNA synthesis. Intronic reads derived from pre-mRNA would align to the genome in the same direction as the mRNA (denoted as forward direction). In contrast, intronic reads derived from genomic DNA align in both forward and reverse directions (i.e. same or opposite direction as the mRNA).

To assess the extent of the different sources of intronic reads, we applied the Abacus workflow to two sNuc-RNAseq datasets from human postmortem brain samples generated using the 10x Genomics Chromium protocol (see Methods). It is worth noting that UMIs from forward and reverse intronic reads on average account for 48.3% and 21.0%, respectively, of total UMIs from reads mapped to gene regions (i.e. exonic and intronic regions) (Table 1). Since reverse intronic reads are primarily derived from genomic DNA, assuming an equal amount forward intronic reads are generated from genomic DNA, we can estimate that UMIs from forward intronic reads derived from pre-mRNAs would account for 27.3% of UMIs (i.e. 21.0% subtracted out from 48.3%). This confirms that pre-mRNA is the primary source of the forward intronic reads but that a substantial portion of forward intronic reads can also be derived from genomic DNA and should not be ignored.

**Table 1.** Number of UMIs identified from reads mapped to different gene regions.

|  | Sample1 |  | Sample2 |  | average<br>percentage |
| --- | --- | --- | --- | --- | --- |
|  | count | percentage | count | percentage |  |
| **UMIs from forward**<br>**exonic reads** | 8042686 | 38.2% | 9416659 | 23.1% | 30.6% |
| **UMIs from forward**<br>**intronic reads** | 10842031 | 51.5% | 18391536 | 45.2% | 48.3% |
| **UMIs from reverse**<br>**intronic reads** | 2183563 | 10.4% | 12921398 | 31.7% | 21.0% |

### Use of UMI counts derived from exonic and intronic reads to improve the identification of nuclei-containing droplets

For all droplet-based sNuc-RNAseq techniques, only a small fraction of the oil droplets formed during the sorting process actually contain a nucleus. One important step in the sNuc-RNAseq data analysis is to identify barcodes that correspond to nuclei-containing droplets from those corresponding to empty droplets containing only ambient RNAs in the solution. Nuclei-containing droplets are expected to have higher UMI counts (i.e. more RNA transcripts) than empty droplets.

Here, we investigated the utility of the Abacus-derived intronic and exonic UMI counts to identify barcodes corresponding to nuclei-containing droplets. For each barcode, we calculated the total UMI count derived from forward intronic reads representing pre-mRNA level, and the total UMI count derived from forward exonic reads representing the mRNA level. Interestingly, we observed two distinct groups of barcodes (Figure 2): Group I with slightly higher UMI counts from forward intronic reads than from forward exonic reads (points close to the diagonal line on Figure 2), and Group II with moderate UMI counts from forward exonic reads (ranging from 100 to 1000 UMIs) but low UMI counts from forward intronic reads (ranging from 10 to 100 UMIs). The default CellRanger pipeline identifies barcodes corresponding to nucleus-containing droplets by a single threshold of UMI counts derived from forward exonic reads. As shown in the green box in Figure 2A, barcodes identified by CellRanger do not adequately capture either of the two distinct populations, resulting in the exclusion of a large number of droplet barcodes. Unlike CellRanger, Abacus filters barcodes by UMI counts from both forward exonic reads and forward intronic reads and recovers most of Group I barcodes as nucleus-containing droplets (points in the red box in Figure 2A). We applied clustering and tSNE visualization on the nuclei identified by Abacus using gene expression values quantified by UMI counts derived from forward exonic reads. Interestingly, these ~1594 barcodes (37% of total barcodes) omitted by CellRanger but recovered by Abacus (point in the red box but outside of the green box in Figure 2A) are grouped into several cell type clusters on the tSNE plot, corresponding to several non-neuronal cell types (e.g. oligodendrocytes, microglia, astrocytes) as well as some neuronal subtypes in the brain. In fact, for the non-neuronal cell types, the CellRanger pipeline has omitted the majority of the nuclei barcodes (89% of the oligodendrocytes, 71% microglia, 75% astrocytes). As a comparison, when including the barcodes from Group II, they form a separate cluster, not expressing any distinct cell type markers, suggesting that Group II barcodes might represent either empty droplets containing ambient RNA or cell debris that survived the nuclear preparation.

**Figure 2.**
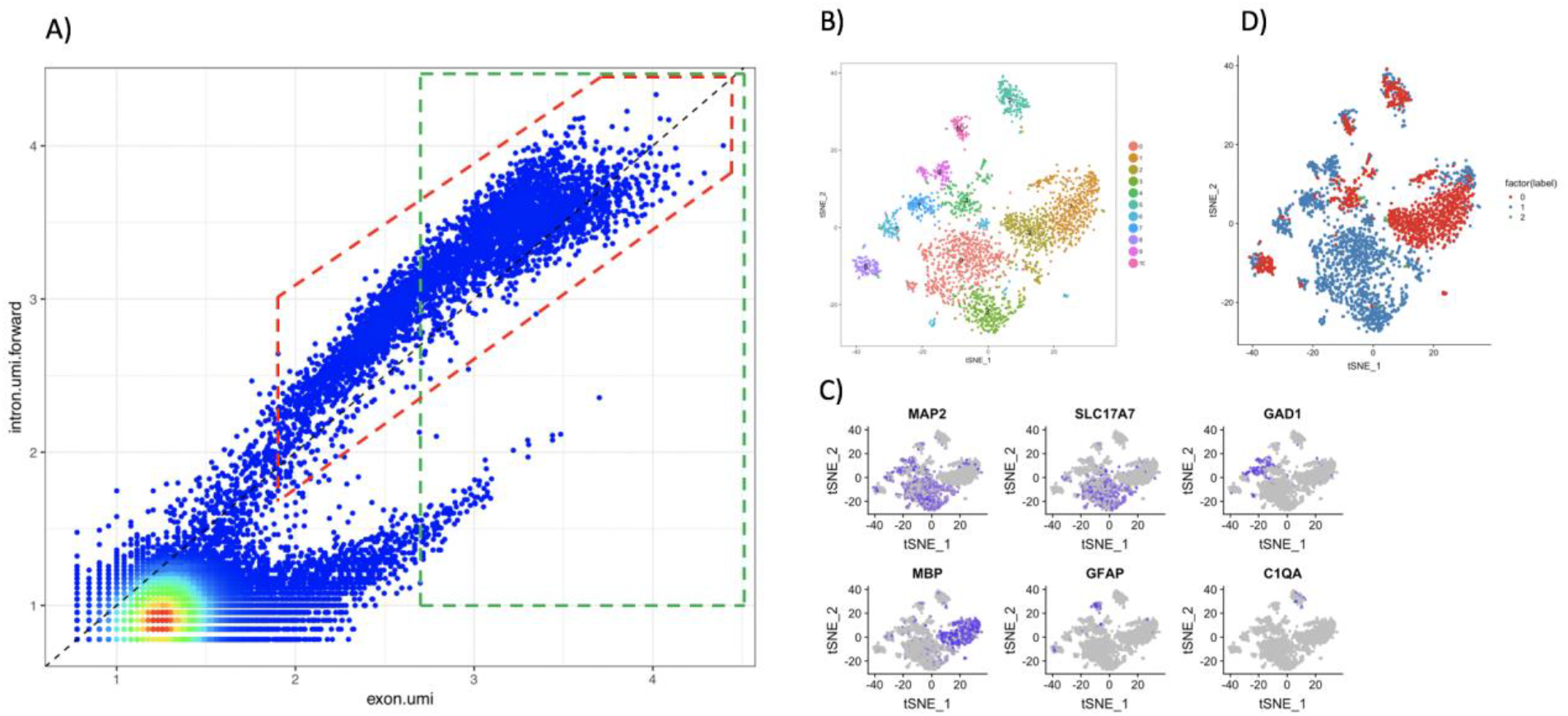
Abacus improves the identification of nuclei from various brain cell types by using UMIs derived from forward intronic and exonic reads. A) Abacus identifies distinct populations of droplets marked by different ratios of forward intronic UMI and exonic UMI counts. The red and green boxes show the final barcodes reported by Abacus and CellRanger respectively. B) Clustering and tSNE visualization of nuclei identified by Abacus. C) Cell type-specific gene markers represented in the distinct clusters (MAP2 = neurons, MBP = oligodendrocyte lineage, SLC17A7 = excitatory neurons, GAD1 = inhibitory neurons, GFAP = astrocytes, C1QA = microglia). D) Nuclei uniquely identified by Abacus belong to diverse brain cell types. Blue dots represent nuclei identified by CellRanger and Abacus, whereas red dots represent nuclei uniquely identified by Abacus. For the non-neuronal cell types, the CellRanger pipeline has omitted the majority of the nuclei barcodes (89% of the oligodendrocytes, 71% microglia, 75% astrocytes; see text).

### Use of UMI counts derived from forward intronic reads for clustering analysis

Even though forward intronic reads could originate from either pre-mRNA or genomic DNA, for abundantly expressed genes, there are far more copies of pre-mRNA transcripts than genomic DNA in the nucleus and, therefore, intronic reads would be predominantly derived from pre-mRNAs and genuinely reflect the gene expression levels. To test this hypothesis, we applied tSNE clustering analysis using the gene expression values quantified by UMIs derived only from forward intronic reads. As expected, clusters corresponding to major brain cell types including excitatory neurons, inhibitory neurons, oligodendrocytes, and astrocytes could be identified. A microglia cell cluster was not detected probably due to the smaller number of microglia cells in the tissue. As Abacus provides separate gene expression quantifications from forward intronic and exonic reads, we could perform cluster analysis jointly with expression values derived from both intronic and exonic reads simply by appending the two feature expression tables. Here, UMI counts derived from exonic reads and intronic reads for the same gene are kept as separate features and contribute to the clustering independently. Joint analysis provides greater cluster refinement as indicated by the increased number of clusters identified at the same level of cluster resolution (Figure 3, 13 clusters identified instead of 10 from clustering using UMI counts derived from exonic reads alone). Taken together, our results strongly suggest that forward intronic reads reflect gene expression levels of the transcriptome and can contribute additional useful information for clustering analysis.

**Figure 3.**
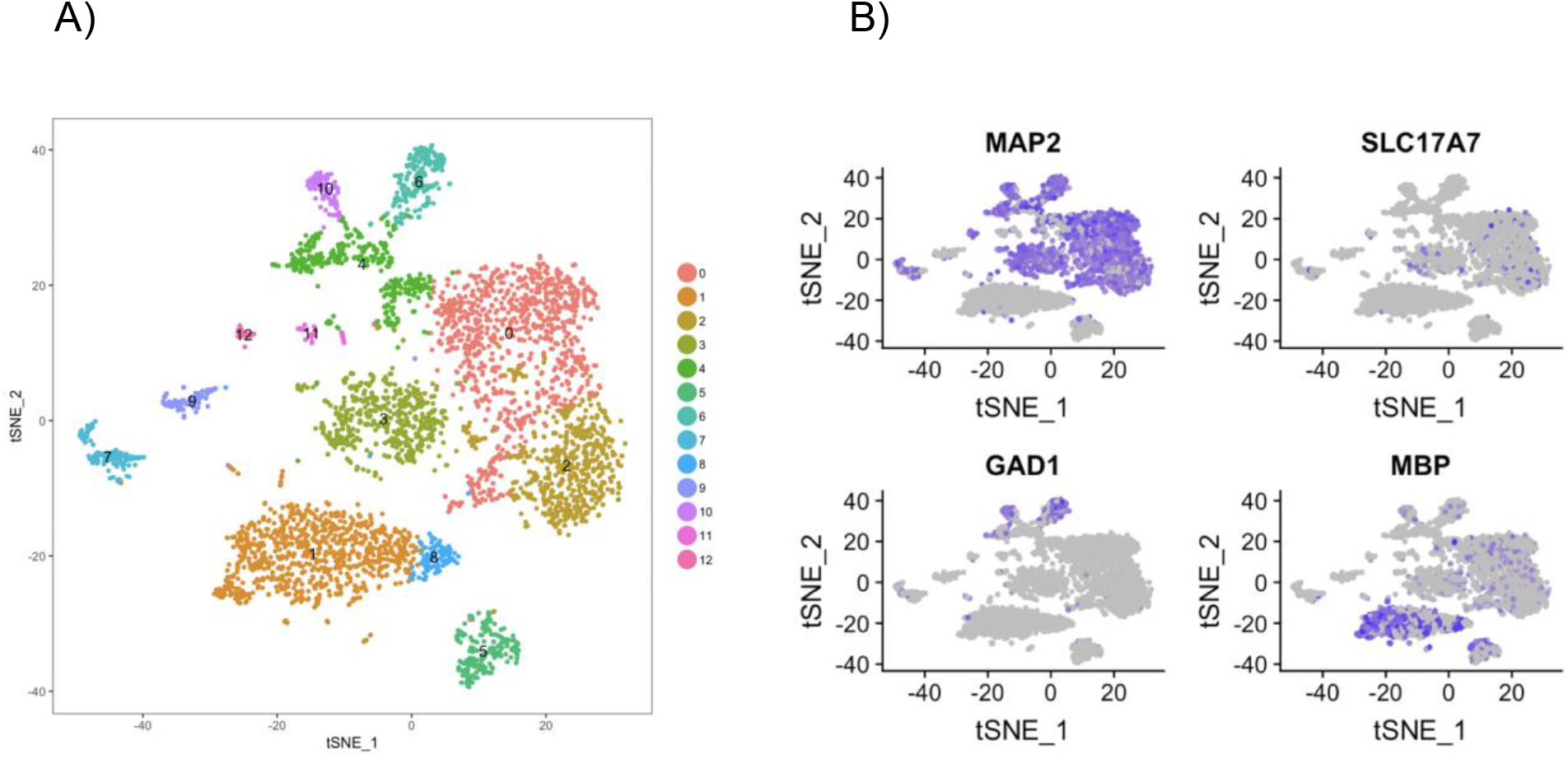
UMIs derived from forward intronic reads reflect gene expression levels of the transcriptome. A) Clustering and tSNE visualization using UMI counts from forward intronic reads. B) Expression of cell-type specific marker genes in the different clusters (MAP2 = neurons, MBP = oligodendrocyte lineage, SLC17A7 = excitatory neurons, GAD1 = inhibitory neurons) demonstrate that clustering based on forward intronic reads maintains clustering by cell type, suggesting a reflection of true gene expression levels.

### UMI counts derived from forward intronic reads cannot be used to quantify gene expression of low abundance genes

Since low abundance genes have very few copies of pre-mRNA in the cell, reads mapped to the intronic regions of these genes would be primarily originated from mis-priming to genomic DNA as opposed to pre-mRNA. Indeed, this is evident from the directionality of the reads. As an example, for the gene FBXW8, a lowly expressed gene as indicated by the few reads mapped at the 3’UTR, intronic reads are mapped to both forward and reverse directions of the FBXW8 gene on the genome (Figure 4B). In comparison, for gene MAP2, a highly expressed gene, intronic reads are predominantly mapped to the forward direction of the MAP2 gene (Figure 4A). Transcriptome-wide analysis reveals that 74% of the genes have more than 3 times the intronic reads mapped to the forward direction over the reverse direction (Figure 4C). This suggests that for a substantial portion of genes (~26%) showing a higher proportion of intronic reads mapped to the reverse direction as compared to the forward direction, the intronic reads may not accurately represent true gene expression levels due to mis-priming to genomic DNA.

**Figure 4.**
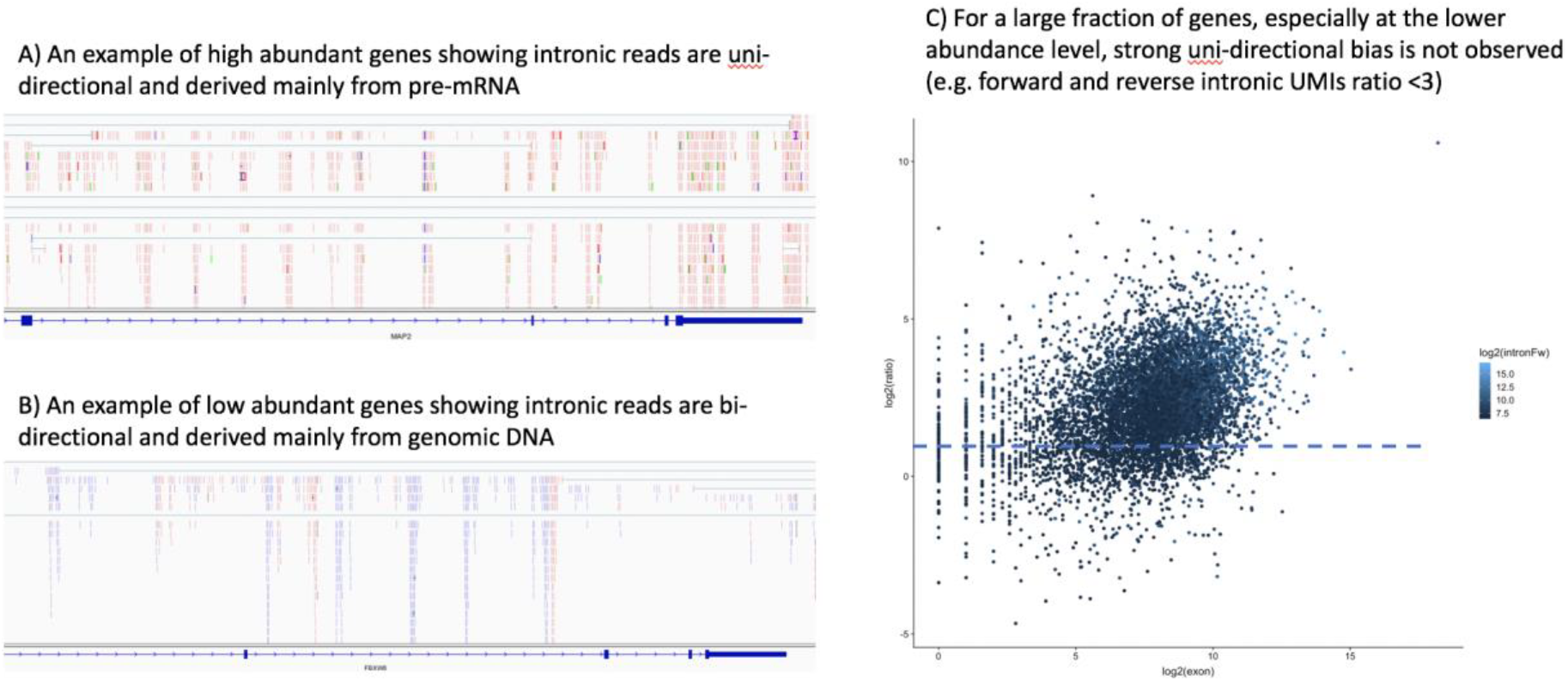
By reporting UMI counts derived from forward, reverse intronic and exonic reads separately, Abacus provides an accurate and flexible representation of gene expression. A) An example of reads mapped to a highly-expressed gene, MAP2, showing that intronic reads map mainly in the forward direction (red) as compared to the reverse direction (blue), and thus are mainly derived from pre-mRNA. B) An example of reads mapped to a lowly-expressed gene, FBXW8, demonstrating a higher proportion of UMI counts in the reverse direction (blue) as compared to the forward direction (red). C) Quantification of the number of UMI counts mapping to exons (x-axis) and the ratio of intronic reads mapping in the forward vs. reverse direction (y-axis). Some genes, especially lowly-expressed genes (i.e. those with lower exonic UMI counts), show a ratio of intronic forward/reverse UMI counts <3, suggesting high reverse intronic read mapping indicative of mis-priming to genomic DNA.

## Discussion

sNuc-RNAseq is often limited by the number of transcripts that can be captured and counted per nucleus, as there are substantially fewer mRNAs in the nucleus than the cytosol. By providing the flexibility to generate UMI counts from reads mapped to different regions of the genome in different orientations, Abacus allows users to make effective use of intronic reads to quantify pre-mRNA transcripts and improves the overall transcript capturing in sNuc-RNAseq studies. By jointly analyzing UMI counts from forward intronic and exonic reads, our analysis revealed distinct subgroups of droplet barcodes. We demonstrated here that, by including UMI counts from forward intronic and exonic reads, Abacus can be used to identify nucleus-containing droplets with better sensitivity and specificity, and substantially improve nucleus barcode calls. This functionality could be particularly useful for sNuc-RNAseq studies of tissues that contain a mixture of cell types with differing RNA content in the nucleus. In our study of human postmortem brain samples, Abacus detected substantially more nuclei of non-neuronal cell types that apparently have fewer UMI counts than neurons.

For gene expression quantification, unlike other tools that combine UMI counts from intronic and exonic reads, Abacus provides users the flexibility to treat UMI counts from forward intronic and exonic reads as separate features. This is particularly useful for analyzing the expression of genes with low abundance, as intronic reads mostly originate from mis-priming of oligo-dT primers to genomic DNA instead of pre-mRNA. For these low abundance genes, tools combining UMI counts from intronic and exonic reads would mis-represent the true expression levels and reduce the sensitivity and specificity in detecting gene expression changes. With Abacus, users can choose to use the different UMI counts depending on the application. For differential gene expression analysis, users can choose to use only UMI counts from exonic reads that provide more accurate representation of low abundance genes. On the other hand, for cluster analysis, UMI counts from both intronic and exonic reads can be used, as the cluster identities are mainly driven by the abundant genes.

## Conclusions

Intronic reads can originate from both pre-mRNAs and genomic DNA, and provide useful features for gene expression analysis in sNuc-RNAseq studies. Abacus is a flexible UMI counter that summarizes intronic and exonic reads from both forward and reverse strands, enabling use of these different UMI counts depending on the analysis context. Joint utilization of UMI counts from forward intronic and exonic reads can improve the identification of nuclei barcodes, especially for cell types with lower transcript numbers, and can increase the refinement of cluster analysis. For genes with low abundance, our analysis indicates that only UMI counts from exonic reads should be used and not combined with UMI counts from intronic reads for gene expression quantification.

## Availability and requirements

Project name: Abacus

Project home page:

Operating system(s): Platform independent

Programming language: Java

Other requirements: Java Runtime Environment (JRE) 8 (64 bit), 4 GB RAM and 1 CPUs

License: GNU GPLv3

Any restrictions to use by non-academics: None

## Acknowledgments/ Disclosures

The design, study conduct, and financial support for the study were provided by AbbVie. AbbVie participated in the interpretation of data, review and approval of the publication. Maya Woodbury, Astrid Wachter, Lauren Gibilisco, Markus Kummer, and Knut Biber are current employees of AbbVie. Simon Xi is a former employee of Abbvie. The authors would like to thank the Alzheimer’s disease research unit at the MassGeneral Institute for Neurodegenerative Disease (MIND) for providing brain tissues and critical feedback on nuclei experiments.

## Notes

### Competing Interest Statement

The authors have declared no competing interest.

